# Free ferrous ions sustain activity of mammalian stearoyl-CoA desaturase-1

**DOI:** 10.1101/2023.03.17.533000

**Authors:** Jiemin Shen, Gang Wu, Brad S. Pierce, Ah-Lim Tsai, Ming Zhou

## Abstract

Mammalian stearoyl-CoA desaturase-1 (SCD1) introduces a double-bond to a saturated long-chain fatty acid and the reaction is catalyzed by a diiron center, which is well-coordinated by conserved histidine residues and is thought to remain with enzyme. However, we find that SCD1 progressively loses its activity during catalysis and becomes fully inactive after nine turnovers. Further studies show that the inactivation of SCD1 is due to the loss of an iron (Fe) ion in the diiron center, and that the addition of free ferrous ions (Fe^2+^) sustains the enzymatic activity. Using SCD1 labeled with Fe isotope, we further show that free Fe^2+^ is incorporated into the diiron center only during catalysis. We also discover that the diiron center in SCD1 has prominent electron paramagnetic resonance signals in its diferric state, indicative of distinct coupling between the two ferric ions. These results reveal that the diiron center in SCD1 is structurally dynamic during catalysis and that labile Fe^2+^ in cells could regulate SCD1 activity, and hence lipid metabolism.

## Introduction

Stearoyl-CoA desaturases (SCDs) are redox-active diiron enzymes embedded in the membrane of endoplasmic reticulum (ER) (Strittmatter et al. 1974; Sperling et al. 2003; Paton and Ntambi 2009). SCDs catalyze the conversion of saturated fatty acids (SFAs) to monounsaturated fatty acids (MUFAs) (Paton and Ntambi 2009). Humans have two SCD isoforms (SCD1 and SCD5) (Wu et al. 2013), while mouse has four (SCD1-4) that are the co-orthologs of human SCD1 (Evans et al. 2008). Studies have shown that mice with SCD1 knocked out are resistant to high-fat diets (Ntambi et al. 2002; Gutiérrez-Juárez et al. 2006), and that SCD1 activity is crucial for maintaining the balance between fat consumption and accumulation (Flowers and Ntambi 2008). Inhibition of SCD1 has been pursued to treat metabolic diseases such as obesity and diabetes (Ntambi et al. 2002; Gutiérrez-Juárez et al. 2006; Xin et al. 2008; Paton and Ntambi 2009; Brown and Rudel 2010; Oballa et al. 2011; Zhang, Dales, and Winther 2014; Sun, Zhang, Raina, et al. 2014; Sun, Zhang, Kodumuru, et al. 2014; Aljohani, Syed, and Ntambi 2017). SCD-mediated desaturation is a major route for the *de novo* synthesis of MUFAs, which is essential for cell survival (Fritz et al. 2010; Peck et al. 2016; Vriens et al. 2019; Oatman et al. 2021). Elevated expression of SCD1 has been observed in various types of cancer cells (Peck and Schulze 2016), and the high level of MUFA production by SCD1 protects cancer cells against ferroptosis (Magtanong et al. 2019; Tesfay et al. 2019; Luis et al. 2021). Preclinical studies of SCD1 inhibitors have shown benefits for the treatment of cancers (Yahagi et al. 2005; Scaglia and Igal 2008; Scaglia, Chisholm, and Igal 2009; Ackerman and Simon 2014; Theodoropoulos et al. 2016; Savino et al. 2020; Oatman et al. 2021). SCD1 is also a validated target for reversing the pathology in neurodegenerative diseases, such as Parkinson’s and Alzheimer’s diseases (Vincent et al. 2018; Fanning et al. 2019; Nuber et al. 2021; Hamilton et al. 2022).

SCDs have a diiron center that undergoes oxidation during the catalysis (Paton and Ntambi 2009), and is recovered by reducing equivalents delivered by a heme protein cytochrome b_5_ (cyt b_5_) (Vergeres and Waskell 1995). Cyt b_5_ receives electrons from cytochrome b_5_ reductase (b_5_R), which obtains electrons from nicotinamide adenine dinucleotide phosphate (NADPH) via a bound flavin adenine dinucleotide (FAD) cofactor (Spatz and Strittmatter 1973; Rogers and Strittmatter 1975; Iyanagi 1977; Iyanagi, Watanabe, and Anan 1984b; Yamada et al. 2013). The three proteins, b_5_R, cyt b_5_, and SCD1, form an ER-resident electron transfer chain that sustains the desaturation reaction with NADH oxidation and O_2_ reduction (Paton and Ntambi 2009) (Figure 1a). SCDs have four transmembrane (TM) helices (Man et al. 2006; Bai et al. 2015; Wang et al. 2015), while both b_5_R and cyt b_5_ have a single TM helix (Vergeres and Waskell 1995; Yamada et al. 2013). In a recent study, we found that the three proteins form a stable ternary complex mediated mainly by the TM helices, and that the formation of a stable ternary complex accelerates electron transfer (Shen et al. 2022).

**Figure 1.**
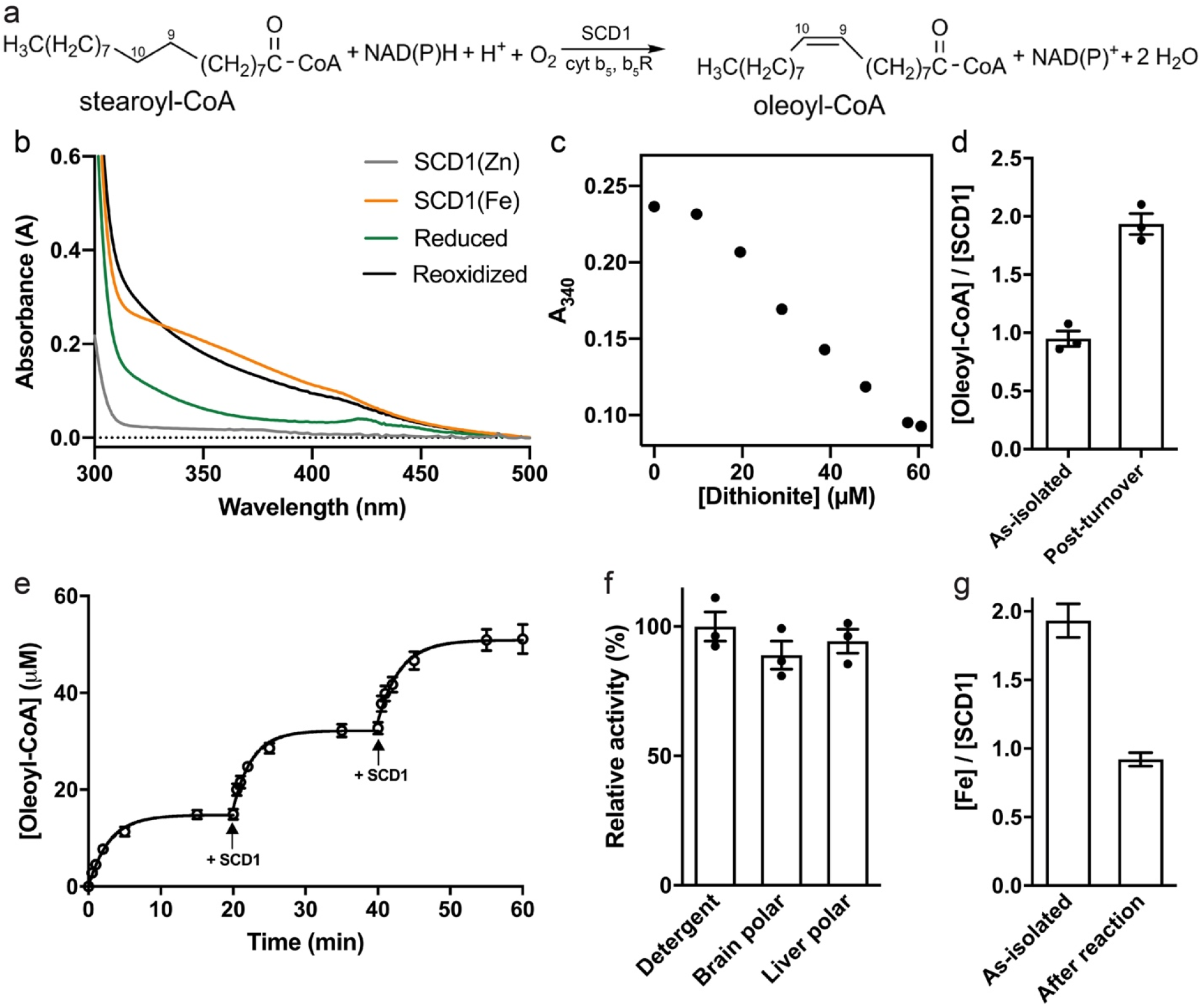
Single turnover and continuous turnover reaction of SCD1. (**a**) Overall reaction scheme of the biological desaturation by SCD1. The Δ9 and Δ10 carbons on the acyl chain are labeled. (**b**) UV/Vis spectra of Zn-containing (gray), Fe-containing (orange), dithionite-reduced (green) and reoxidized (black) SCD1. (**c**) Anaerobic titration of dithionite into 50 μM as-isolated SCD1. (**d**) Molar ratios of oleoyl-CoA to SCD1 in samples before and after chemical reduction and reoxidation in the presence of substrate stearoyl-CoA. (**e**) Time course of oleoyl-CoA production by SCD1 in the biological pathway with cyt b_5_ and b_5_R. The same amount of SCD1 was added at time points indicated by black arrows. (**f**) Activities of SCD1 in detergent or in liposomes prepared with brain polar or liver polar lipids. (**g**) Fe content analysis of SCD1 before and after reaction by ICP-MS. The molar ratios of Fe to protein are shown. In this paper, error bars represent the standard error of the mean from three repeats.

Initial structures of SCD1s have a diiron center that was mis-incorporated with two zinc ions (Zn^2+^), an artifact likely stemmed from overexpression of the proteins in insect cells (Bai et al. 2015; Wang et al. 2015). In a follow-up study, we developed a protocol to produce mammalian SCD1 with >90% iron (Fe) occupancy and determined its crystal structure (Shen et al. 2020). These studies (Bai et al. 2015; Wang et al. 2015; Shen et al. 2020) defined a unique diiron center coordinated entirely by histidine residues: two Fe ions (Fe1 and Fe2) are coordinated by the imidazole side chain of nine histidine residues, five for Fe1, four for Fe2; the Fe···Fe distance is 6.4 Å. This configuration is different from previously resolved diiron centers (Lindqvist et al. 1996; Högbom et al. 2002; Sazinsky and Lippard 2006; Jasniewski and Que 2018), which contain carboxylate ligands with one or two carboxylates forming bidentate bridge(s) between the two Fe ions, restricting the Fe···Fe distance to less than 4 Å. The distinct configuration of the diiron center in SCD1 is representative of many membrane-bound desaturases and hydroxylases (Shanklin, Whittle, and Fox 1994; Sazinsky and Lippard 2006; Zhu et al. 2015), but its reaction mechanism remains unclear (Jasniewski and Que 2018). In this study, we report the catalytic inactivation of the diiron center in SCD1 and the regulation of SCD1 activity by exogenous ferrous ions (Fe^2+^).

## Results

### Fast inactivation of SCD1

In a previous study, we purified Fe-loaded mouse SCD1 from HEK cells and verified its Fe content in two ways (Shen et al. 2020). First, we measured Fe content using inductively coupled plasma mass spectrometry (ICP-MS), and found that the molar ratio of Fe:SCD1 is ∼2:1. Second, we verified the presence and location of the diiron center in the crystal structure by the anomalous X-ray dispersion signals of Fe. Here, we further characterized the diiron center. We chemically reduced the diiron center in the purified SCD1 by anaerobic stoichiometric titration with dithionite, and monitored the decrease of absorbance of diferric cluster (Fe(III)/Fe(III)) at 340 nm (A_340_) (Methods and Figure 1b). We found that ∼2 reducing equivalents are required to fully reduce the diiron center in the purified SCD1 (Figure 1c), which further confirms the 2:1 [Fe]:[SCD1] ratio, and indicates that the resting diiron center in purified SCD1 is in the Fe(III)/Fe(III) state. We then show that SCD1 reduced by dithionate is enzymatically active, and produces one equivalent of product in the presence of stearoyl-CoA and O_2_, indicative of a single turnover reaction (Figure 1d).

We proceed to examine SCD1 activity under conditions that enable multiple turnovers. In the presence of purified mouse cyt b_5_ and b_5_R and sufficient amount of substrate stearoyl-CoA and NADH (1 mM), the formation of product oleoyl-CoA levels off with an average total turnover number (TTN) of 8.5 ± 0.3 (Figure 1e). However, when fresh SCD1 is added to the reaction mixture, we observe additional product formation with almost identical initial rate and TTN to the initial round (Figure 1e). And this process can be repeated with the same outcome. There seems to be a linear relationship between the amount of SCD1 and the yield of oleoyl-CoA, whether the enzyme is added in increments or all at the beginning (Shen et al. 2020). Thus, the loss of enzymatic activity is not due to the exhaustion of reducing equivalents or substrates, nor is it due to product inhibition. We conclude that SCD1 becomes progressively self-inactivated after each turnover and we surmise that this is due to the loss of Fe in SCD1.

To alleviate concern that detergent-solubilized SCD1 becomes unstable during the reaction, we conduct the following two experiments. First, we reconstitute purified SCD1 into liposomes for the enzymatic assay, and we observe a similar loss of activity as the reaction progresses (Figure 1f). We also notice that in an earlier study of SCD1 from crude extract of liver microsomes, self-inactivation was obvious with a similarly TTN (Joshi, Wilson, and Wakil 1977). Second, we purified the inactivated SCD1 from the reaction mixtures by size-exclusion chromatography (SEC). A monodispersed peak in SEC (Figure 1—figure supplement 1a) and a single band in SDS-PAGE (Figure 1—figure supplement 1b) indicate that the inactivated SCD1 is biochemically stable as it was before the reaction. We conclude that the inactivation of SCD1 is not due to protein aggregation or loss of its lipidic environment.

### Loss of SCD1 activity in the peroxide-shunt pathway

Similar to cytochrome P450, diiron enzymes can react with hydrogen peroxide (H_2_O_2_) to drive enzymatic turnover in the absence of an electron transfer chain, which is referred to as the “peroxide shunt” pathway (Bailey and Fox 2009; Jasniewski and Que 2018). This pathway has not been reported in SCD1. We find that SCD1 can utilize the peroxide-shunt pathway (Figure 1—figure supplement 2a), although the rate of substrate formation is (Figure 1—figure supplement 2b). The *K*_M_ for H_2_O_2_ is 18.4 ± 2.3 mM and the *k*_cat_ is 0.84 ± 0.03 min^-1^ (Figure 1— figure supplement 2c), which is ∼3.3-fold slower than the biological pathway. SCD1 also displays progressive loss of activity with a TTN of 6.8 ± 0.3, but over a longer period of time (∼30 min) (Figure 1—figure supplement 2b). Although the peroxide-shunt pathway may not be biologically relevant, it incurs a similar loss of enzymatic activity. This leads us to hypothesize that SCD1 activity is vulnerable to self-inactivation during the oxidation of the diiron center.

We also examined the kinetics of the peroxide-shunt pathway in SCD1 by following the UV-Vis spectra of the diiron center after rapid mixing. The time course can be deconvoluted into two major phases (1 and 2) with transition rates of 9.7 s^-1^ and 0.17 s^-1^ (Figure 1—figure supplement 2d), which reflect the reaction of the diiron center with H_2_O_2_. These rates are much faster than the *k*_cat_ of oleoyl-CoA production, suggesting that the H_2_O_2_ activation of the diiron center is not the rate-limiting step.

### Loss of one Fe during catalysis

We measured the amount of Fe in the inactivated SCD1 by ICP-MS, and found that the [Fe]:[protein] ratio drops from ∼2:1 to ∼1:1 (Figure 1g). It is more likely that most of the SCD1 has lost a single Fe ion than the alternative interpretation that there is a mixture of SCD1 with zero, one, or two Fe ions. We will further examine the stoichiometry in the Fe exchange experiments and in electron paramagnetic resonance (EPR) analyses described later in the manuscript. We also found that the diiron center in the purified SCD1 is resistant to chelation by ethylenediaminetetraacetic acid (EDTA) (Methods). We speculate that the loss of Fe occurs only during the catalytic cycle, likely when the diiron center experiences a higher oxidation state.

### Exogenous Fe^2+^ enhances the activity of SCD1

We then examined the activity of SCD1 in the presence of free Fe^2+^ in the solution and found that Fe^2+^ can sustain the reaction as evidenced by significantly higher amount of substrate conversion (Figure 2a). Other common transition metal ions were also tested, but none of them significantly changes the TTN of SCD1, including ferric ion (Fe^3+^) (Figure 2b). Ascorbate, added to protect Fe^2+^ from oxidation in solution, does not enhance the activity of TTN by itself (Figure 2b). The enhancement by Fe^2+^ is concentration-dependent with a half maximal effective concentration (EC_50_) of 6.5 (4.5–9.4) μM (95% confidence interval in parentheses) (Figure 2c). Further, we found that addition of free Fe^2+^ prior to the start of the reaction does not increase the initial rate of reaction (Figure 2d, inset), but only increases the TTN in the reaction system. After SCD1 is inactivated, however, addition of Fe^2+^ does not recover the enzymatic activity (Figure 2e), indicating that the protection of enzymatic activity by free Fe^2+^ occurs during the catalytic cycle.

**Figure 2.**
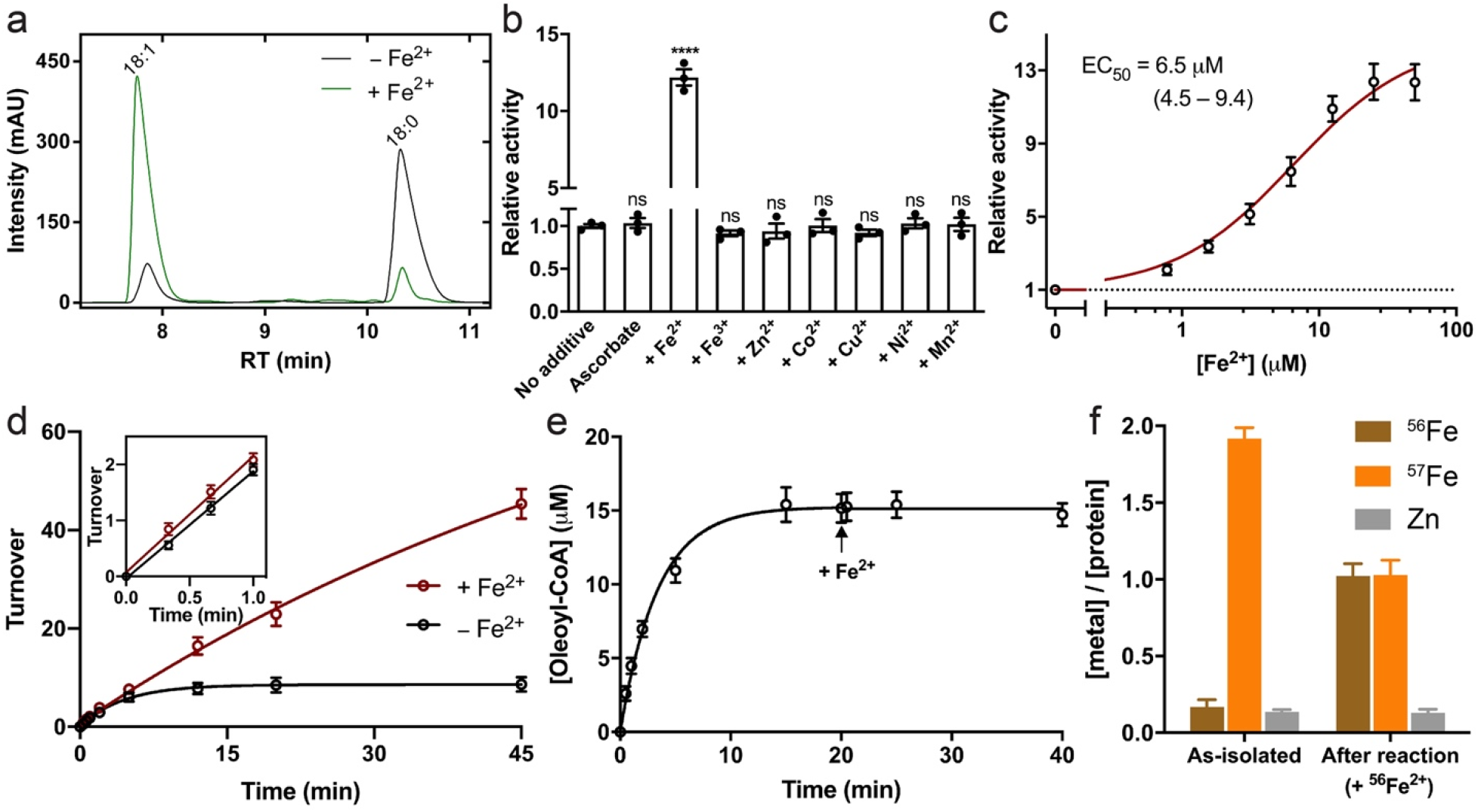
Enhanced activity of SCD1 in the presence of exogenous Fe^2+^. (**a**) HPLC profiles of acyl-CoA conversions in the absence (black) or presence (green) of Fe^2+^. Peaks for the product (18:1) and substrate (18:0) are labeled. (**b**) Activities of SCD1 in the presence of exogenous transition metal ions. Statistical significances were analyzed with one-way analysis of variance (ANOVA) followed by Dunnett’s test for multiple comparisons. ns, not significant; ****, *p* < 0.0001. (**c**) Concentration-dependent potentiation of SCD1 activity by exogenous Fe^2+^. The EC_50_ value was calculated from the fit (red line) to an excitatory dose-response equation. 1 mM ascorbate-Na was added together with Fe^2+^. (**d**) Time courses of oleoyl-CoA production in the presence (red) or absence (black) of exogenous Fe^2+^. The inset shows that the initial rates are not significantly different in the two conditions. (**e**) No additional product formation after the addition of Fe^2+^ (black arrow) to inactivated SCD1. No exogenous Fe^2+^ was present prior to the start of reaction. (**f**) Metal content analysis of ^57^Fe-enriched SCD1 before and after reaction. ^56^Fe^2+^ was added prior to the start of reaction. SCD1 was separated for analysis after 30 min reaction.

### Fe exchange occurs during catalysis

To further understand how free Fe^2+^ helps prevent the inactivation of SCD1, we prepared SCD1 enriched with ^57^Fe to track changes of Fe in the diiron center (Methods). We first expressed and purified ^57^Fe-enriched SCD1 (Methods), and recovered the ^57^Fe-enriched SCD1 after ∼30 min reaction in the presence of free ^56^Fe^2+^ in the solution. ICP-MS shows that while the total [Fe]:[protein] ratio is still close to 2:1 (Figure 2f), the isotope composition changes dramatically: the endogenous [^57^Fe] in SCD1 drops by ∼50%, and the exogenous [^56^Fe] becomes comparable to [^57^Fe], making the [^56^Fe]:[^57^Fe]:[protein] ratio close to unity (Figure 2f). As expected, incubation of ^56^Fe^2+^ with ^57^Fe-enriched SCD1 for ∼1 h without initiating the catalysis does not allow the incorporation of ^56^Fe^2+^ into the diiron center (Figure 2—figure supplement 1), indicative of a stable diiron center at the resting state. We conclude that free (exogenous) Fe^2+^ is only able to replace the bound (endogenous) Fe during catalysis. Since SCD1 in the absence of free Fe^2+^ is fully inactivated after 30 min while SCD1 in the presence of Fe^2+^ remains fully active after 30 min (Figure 2d), we conclude that the mixed isotope ^57^Fe/^56^Fe center is likely the dominant species in the SCD1 sample after the reaction.

### Unique EPR signatures of resting SCD1

Since the diiron center in the purified SCD1 is in the Fe(III)/Fe(III) state, we proceed to examine the diiron center using EPR spectroscopy. Previous studies of the diiron centers in several enzymes show that they are EPR-silent in the Fe(III)/Fe(III) state due to the antiferromagnetic (AF) coupling of two Fe(III), likely via bridging ligand(s) (Jasniewski and Que 2018). However, due to the long Fe···Fe separation (6.4 Å) and lack of a bridging ligands in SCD1, we consistently obtain EPR signals from SCD1. SCD1 has a sharp asymmetric species at *g* ∼4.3 (Figure 3a, I) with a half-saturation power (P_1/2_) of ∼141 mW (Figure 3b), which is typical of highly rhombic high-spin Fe(III) EPR. Interestingly, SCD1 also has a prominent broad species with a linewidth of ∼70 mT centered at nominal *g* ∼2.0 (Figure 3a, II). Species II relaxes extremely efficiently with a P_1/2_ higher than 200 mW (Figure 3b). Crucially, the temperature dependence of II does not follow Curie Law behavior. Instead, the temperature-normalized signal intensity (S×T) increases with temperature indicating that an excited “EPR-active” spin manifold is being populated. This observation is inconsistent with the behavior expected for known (*S* = 1/2) mixed valent Fe(II)/Fe(III) or Fe(III)/Fe(IV) clusters. While much more work is needed to assign this specific EPR transition, the observed temperature dependence of II can be modeled as a transition within an excited *S* = 1 manifold assuming weak (*J* = −7.5 cm^-1^) antiferromagnetic coupling of two high spin Fe(III) sites. If correct, the magnitude of this AF coupling would be the lowest reported among enzymatic Fe(III)/Fe(III) clusters (Jasniewski and Que 2018). Additional work and quantitative simulations are needed to fully validate this assignment.

**Figure 3.**
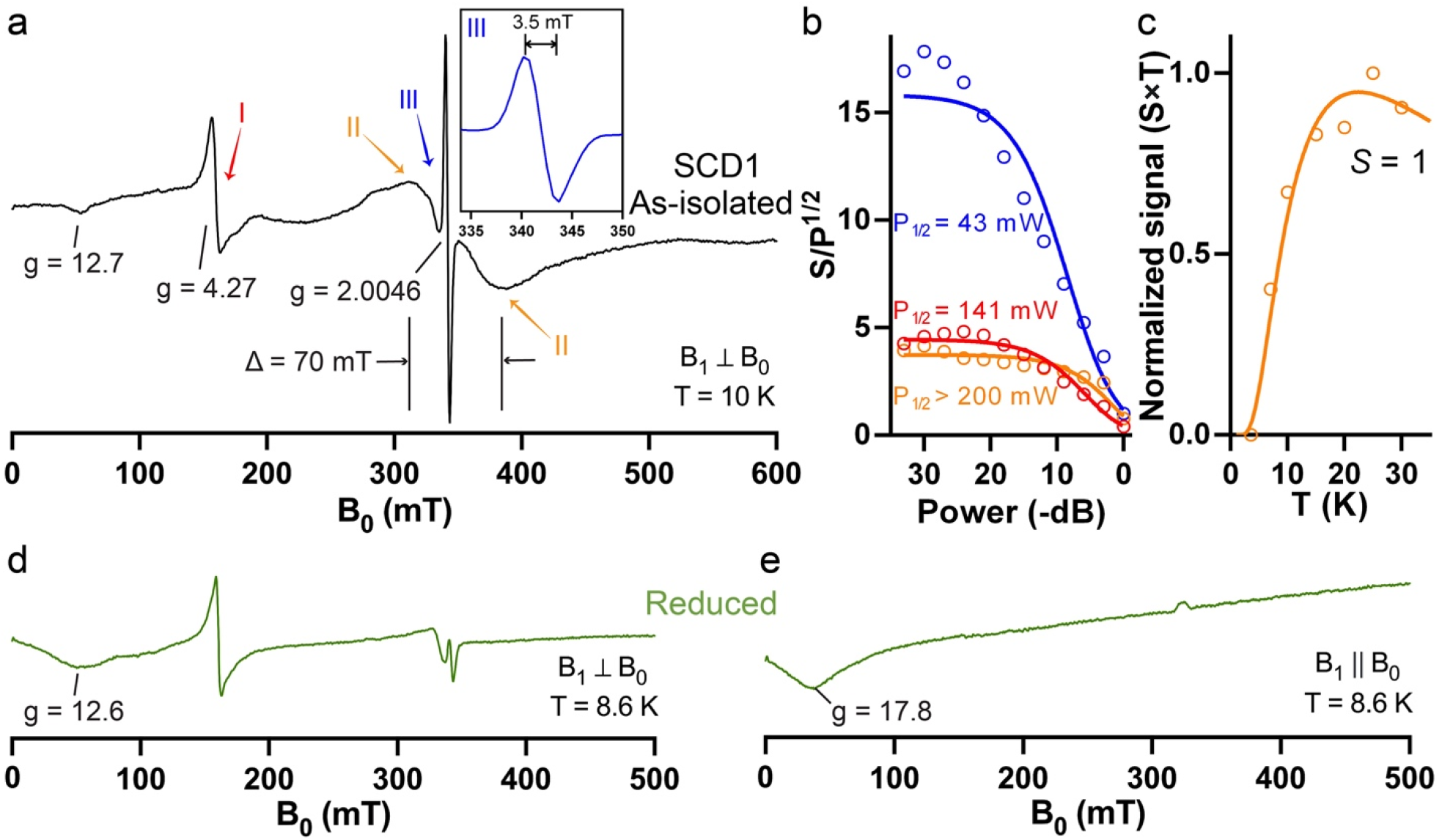
EPR spectroscopy of the diiron center in SCD1. (**a**) As-isolated SCD1 (1 mM) at 10 K shows strong EPR signals around *g* = 4–6 (I), a broad symmetric signal centered at *g* ∼2 with a peak-to-trough span (Δ) = 70 mT (II), and a sharp radical-like signal (III). Inset: Narrow field window of EPR species III (blue). (**b**) Power dependence of EPR species I (red), II (orange), and III (blue) at 10 K. (**c**) The temperature dependence of the normalized signal of species II from *S* = 1 (orange circles) and the calculated Boltzmann curve (orange line). EPR spectrum of SCD1 after photoreduction (green) in perpendicular mode (**d**) or in parallel mode (**e**).

Results presented above are consistent with the conclusion that EPR species II originates from a weakly coupled Fe(III)/Fe(III) cluster in resting SCD1 through a mechanism currently under investigation. Significantly, this broad EPR species is not observed in samples prepared from His265Leu SCD1. This SCD1 variant has a crippled Fe2 binding site (see below) and thus cannot produce a diiron cluster.

Apart from the broad II species, multiple batches of SCD1 exhibit a consistent narrow EPR signal centered at *g* ∼2.0 (Figure 3a, III). This feature has a linewidth of ∼3.5 mT with an isotropic lineshape (Figure 3a, inset). The *g*-value and narrow linewidth of the species suggests an organic radical. However, as shown in Figure 3b, the power required for half-saturation (P_1/2_, 43 mW) is significantly higher than typically observed for organic radicals. The fast relaxation observed for species III suggests that it is likely close to another paramagnetic center, presumably the coupled diiron center.

The EPR spectrum of SCD1 has significant changes upon reduction. Photo-reduction that converts the diiron center to the diferrous (Fe(II)/Fe(II)) state largely eliminates EPR signals II and III (Figure 3d). The fully reduced Fe(II)Fe(II) diiron cluster can be observed in the perpendicular mode (9.58 GHz) EPR spectra as a weak dip with the trough centered at an apparent *g*-value of ∼13. This cluster is more readily observed by switching to parallel mode (9.23 GHz) where the Fe(II)Fe(II) cluster exhibits a characteristic signal near *g* ∼18 (Figure 3e).

### Single turnover of SCD1 with cyt b_5_ and b_5_R monitored by EPR

We then examined redox-dependent changes of the EPR signals from SCD1 when reacted with its physiological electron transfer partner cyt b_5_, which is in turn reduced by b_5_R using NADH. To ensure efficient electron transfer, we used a stable ternary complex composed of SCD1, full-length cyt b_5_, and b_5_R reported previously (Shen et al. 2022). The EPR spectrum (at 10 K) of the resting state complex shows distinguishable signatures from low-spin heme Fe(III) of cyt b_5_ at *g* ∼2.17 and *g* ∼3.05 (Guzov et al. 1996), and from SCD1 at *g* ∼4.3, *g* ∼2.1, and *g* ∼2.0 (Figure 3— figure supplement 1a). SCD1 in the ternary complex exhibits similar EPR signatures as the individual SCD1. We then added an equal molar amount of NADH to the complex and followed redox changes by freeze quench. After 10 s, the *g* ∼2.0 signal intensifies dramatically, which has a linewidth of only ∼1.5 mT, consistent with an anionic FAD semiquinone radical (FAD^**-**^**·**) (Iyanagi, Watanabe, and Anan 1984a) in b_5_R. The FAD^**-**^**·** is generated from the rapid hydride transfer from NADH to FAD in b_5_R and subsequent one-electron transfer from FADH_2_ to cyt b_5_ and O_2_ (Figure 3—figure supplement 1d). The disappearance of the heme Fe(III) signals marks the full reduction of cyt b_5_ (Figure 3—figure supplement 1c). Moreover, the *g* ∼2.1 signal from SCD1 starts to decrease, reflecting the fast electron transfer from cyt b_5_ to the diiron center (Figure 3—figure supplement 1b). After another 30 s, the *g* ∼4.3 signal from SCD1, which stays almost unchanged after 10 s reaction, starts to drop (Figure 3—figure supplement 1b), suggesting relatively slower reduction kinetics. While the signal of FAD^**-**^**·** disappears within ∼160s (Figure 3—figure supplement 1d), signals from cyt b_5_ and SCD1 gradually recover over ∼30 min due to slow re-oxidation of the heme and diiron center by O_2_ (Figure 3—figure supplement 1b-c). Compared to the *g* ∼2.1 signal, the *g* ∼4.3 signal has a slower reoxidation rate (Figure 3—figure supplement 1b). These observations support our assignment of the peaks in the EPR spectrum of SCD1, and establish EPR spectroscopy as a powerful approach for further investigation of the mechanism of the diiron center.

### Loss of Fe2 in SCD1

EPR spectrum provides an independent way of assessing the loss of Fe in SCD1. EPR spectra collected on inactivated SCD1 exhibit multiple signals which can be attributed to high-spin Fe(III). In addition to the sharp *g* ∼4.3 signal attributed to a transition within the middle doublet (m_s_ = ±3/2) of the high-spin state, weaker signals are also observed at *g*-values ranging from 5 to 9 (Figure 4a). Significantly, the broad species II and the sharp species III found in the fully active SCD1 are not observed in the inactivated SCD1. These observations, combined with the ICP-MS result, support our hypothesis that the inactivated SCD1 is predominantly bound with a single Fe in the diiron center.

**Figure 4.**
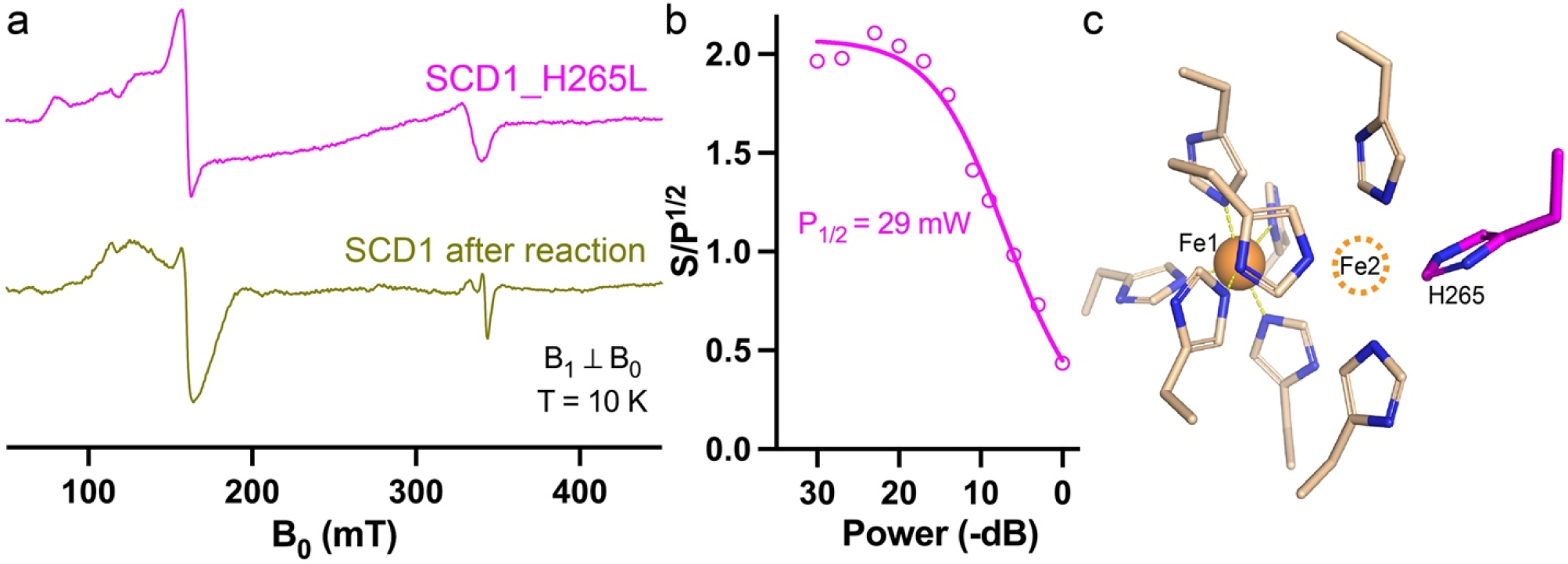
Loss of Fe detected by EPR spectroscopy. (**a**) Comparison of EPR spectra of SCD1 with H265L mutation (magenta), 850 μM; SCD1 after catalytic inactivation (yellow), 800 μM. (**b**) Power dependence of EPR signals of H265L at 10 K. (**c**) Structure of the diiron center in SCD1 (PDB ID: 6WF2) showing nine histidine residues in the first coordination sphere. Fe2 is drawn as an orange dashed circle. H265 is highlighted in magenta.

To further identify which Fe is lost from SCD1, we mutated each of the nine Fe-coordinating histidine residues to leucine. However, out of the nine histidine-to-leucine point mutations, only His265Leu that coordinates Fe2 is biochemically stable and has sufficient yield for EPR study (Figure 4—figure supplement 1a). The Fe occupancy of His265Leu SCD1 is slightly less than one iron per SCD1 (Figure 4—figure supplement 1b) and it does not have enzymatic activity. His265Leu SCD1 has EPR signals at *g* 5–9 and ∼4.3 (Figure 4a), but no broad species II and sharp species III were observed in the wild-type (WT) SCD1. Compared with WT SCD1, the saturation behavior (P_1/2_, 29 mW) of His265Leu SCD1variant is much lower (Figure 4b). This is likely attributed to the absence of the adjacent Fe2 ion, as the presence of a nearby paramagnetic center would increase spin relaxation, leading to higher P_1/2_. These features in the EPR spectrum of His265Leu are consistent with a crippled diiron center singly occupied at Fe1 (Figure 4c). Similar EPR signatures of the inactivated SCD1 and the His265Leu SCD1 imply that Fe2 is lost during the catalysis, although we cannot fully exclude the possibility of loss of Fe1 due to lack of EPR spectra from SCD1 singly occupied with Fe2.

## Discussion

In summary, we found that SCD1 catalyzes on average ∼8.5 cycles of the desaturation reaction before fully inactivate, and that the inactivation is caused by the loss of one Fe, most likely Fe2, in the diiron center. The presence of free Fe^2+^ in solution can replace the lost Fe during the enzymatic reaction and thus alleviates or prevents inactivation. However, Fe^2+^ is not able to revive SCD1 once it has already been inactivated. These behaviors were not reported in any other diiron enzymes and seem unique to SCD1 and, by extension, to membrane-bound desaturases that contain a similar diiron center, such as fatty acid desaturase (FADS2) and sphingolipid Δ4-desaturase-1 (DES1) (Figure 4—figure supplement 2).

We notice that the purified SCD1 has a bound oleoyl-CoA (Figure 1d), indicative of a rate-limiting step in product release. It is possible that the loss of one Fe could occur during product release, which requires structural changes. Alternatively, if a highly reactive species is generated in SCD1, for example, Fe(IV)=O^2-^ (Yu and Chen 2019), it may irreversibly damage surrounding residues (Gray and Winkler 2018; Esselborn et al. 2019) such as the histidine residues that coordinate Fe2. The damage may be prevented in the presence of free Fe^2+^. Self-inactivation was observed in prostaglandin H-synthase and the culprit is likely the ferryl heme intermediate rather than the active tyrosyl radical (Wu et al. 2007). In nitric oxide synthase, when either the substrate or the tetrahydrobiopterin cofactor is absent, the P450-like heme can generate superoxide and hydrogen peroxide reactive oxygen species and alter the reaction from nitric oxide synthesis to superoxide (or hydrogen peroxide, or even peroxynitrite) synthesis (Xia et al. 1998; Berka et al. 2004). Without a bridging ligand between the two Fe ions, the increased structural flexibility may enhance such damaging side reactions to inactivate the enzyme when a high oxidation state or peroxo intermediate is formed.

We report the first EPR spectrum of the diiron center in SCD1. Distinct from other well-characterized soluble diiron enzymes, such as ribonucleotide reductase, soluble methane monooxygenase, and acyl-acyl carrier protein (ACP) desaturase, SCD1 in the resting state has a broad EPR species which likely comes from coupled Fe1 and Fe2. The assignment of this EPR signature is supported by its disappearance in the His265Leu mutant and inactivated SCD1 (Figure 4a), both of which contain only one Fe. This EPR signature is also sensitive to changes in the redox state, as observed in chemically reduced SCD1 (Figure 3d–e) as well as in the biological pathway with cyt b_5_ and b_5_R (Figure 3—figure supplement 1). The EPR of SCD1 also shows a radical signal at *g* ∼2.0, which suggests that a high-valent Fe could generate an amino acid radical(s) in its vicinity. We anticipate that refined assignments of the rich EPR signatures of SCD1 will help gain insights into the electronic structure and reactivity of the unique diiron center in the future.

The enhancement of the desaturation activity of SCD1 is dependent on the concentration of free Fe^2+^ in solution. The EC_50_ (6.5 μM) of the Fe^2+^ potentiation seems relevant to typical Fe concentration (∼2 μM) in the cellular labile iron pool (LIP) (Hider and Kong 2011; Cabantchik 2014), which includes Fe^2+^ in complex with chelators such as glutathione and ascorbate. Thus, the LIP could be a regulatory factor of SCD1 activity.

## Methods

### Expression and purification of SCD1 and related proteins

The codon-optimized cDNA of N-terminal truncated mouse SCD1 (Δ2–23) and full-length human SCD5 were cloned into pEGBacMam vectors with a C-terminal enhanced green fluorescent protein (eGFP) tag. Expression was performed through transduction of HEK 293S cells with baculoviruses produced in Sf9 (*Spodoptera frugiperda*) cells following the standard BacMam protocol (Goehring et al. 2014) as previously reported. *FreeStyle 293* media (Invitrogen/Thermo Fisher) were supplemented with apo-transferrin (Athens Research & Technology) and FeCl_3_ (Sigma). 10 mM sodium butyrate (Sigma) was added 1 day after viral transduction and the temperature was lowered from 37 °C to 30 °C. For expression of SCD1-cyt b_5_-b_5_R ternary complex, 0.5 mM δ-aminolevulinic acid (Santa Cruz) and 100 μM riboflavin (Sigma) were added to media to enhance the incorporation of heme and FAD, respectively. Cell membranes were solubilized with 30 mM n-Dodecyl-β-D-Maltopyranoside (DDM, Anatrace). Home-made eGFP nanobody-crosslinked resins (NHS-Activated Sepharose 4 Fast Flow, Cytiva) were used to capture target proteins. EGFP tags were removed after Tobacco Etch Virus (TEV) protease digestion, which also cleaves the interdomain linkers in the ternary complex. Proteins were concentrated with ultrafiltration centrifugal devices of 50-kDa cutoff (Amicon, Millipore). Monodispersed fractions of proteins were collected from size-exclusion columns (Superdex 200 10/300 GL, GE Health Sciences) equilibrated with FPLC buffer (20 mM HEPES, pH 7.5, 150 mM NaCl, 1 mM DDM).

Soluble mouse cyt b_5_ (1–89) and soluble mouse b_5_R (30–301) were expressed in *E. coli* (*Escherichia coli*) BL21(DE3) as previously reported (Shen et al. 2020). Terrific Broth (TB) media were supplemented with δ-aminolevulinic acid or riboflavin. Cobalt-based affinity resins (Talon, Clontech) were used to capture target proteins with an N-terminal His-tag, which was removed after TEV protease digestion.

### Production of ^57^Fe-enriched SCD1

To remove Fe in commercial *FreeStyle 293* media, 20 g of Chelex 100 chelating resins (Bio-Rad) were added per 1 L and incubated under stirring at 4 °C for 5 days. Media were sterile-filtered after the treatment. ^57^FeCl_3_ stock was prepared by dissolving ^57^Fe powder (Isoflex) in 0.1 N HCl. 20 μM of sterile-filtered FeCl_3_ solution together with apo-transferrin were added to the treated media. Other essential divalent metal ions, including Mg^2+^ and Ca^2+^, which were also removed by the chelating resins, were supplemented to the treated media per the previous report on the metal contents of *FreeStyle 293* media (Richardson et al. 2018). The media were pH-adjusted with NaOH or HCl to pH = 7.4. Despite the replenishment of essential metal ions, the treated media do not sustain continuous growth of HEK cells. To minimize cell death, culture media were not exchanged until viral transduction. About 16 h after viral transduction, cells in normal media were pelleted at 800xg and resuspended in the treated media with 10 mM sodium butyrate. Then, the suspended cells were incubated at 30 °C for 2 days before harvest. Purification was conducted as described above for normal SCD1.

### Enzymatic assays of SCD1

Desaturation reactions of SCD1 with soluble cyt b_5_ and b_5_R were performed in conditions similar to those in previous reports (Shen et al. 2020, 2022). SCD1 was mixed with 5x molar excess of soluble cyt b_5_ and b_5_R in FPLC buffer. Stearoyl-CoA (18:0) (Sigma) was used as substrate. For activity assays in liposomes, SCD1 was reconstituted into liver polar extract (Avanti) or brain polar extract lipids (Avanti) following a previous protocol (Shen et al. 2022). When testing the effects of transition metal ions, freshly prepared metal chloride salt stock solutions were added prior to the addition of NADH. Fe^2+^, premixed with a sodium ascorbate stock, was added to achieve a final [ascorbate] of 100 μM and a desired [Fe^2+^]. Reactions were initiated by the addition of 1 mM NADH. For the peroxide-shunt reaction, only SCD1 and substrate stearoyl-CoA were included. H_2_O_2_ (Sigma) was added to start reactions. Aliquots of reaction mixtures were quenched at certain time points. Protein aggregates were pelleted by centrifugation. Supernatants containing acyl-CoAs were analyzed in high-performance liquid chromatography (HPLC). Calibration curves were generated from standard acyl-CoA samples. The initial rates were calculated from the linear fitting of time courses within the first 1 min for biological reactions with cyt b_5_ and b_5_R, or within the first 5 min for peroxide-shunt reactions. Relative activities were based on TTNs of reactions.

### ICP-MS

Protein samples for ICP-MS were collected from SEC with Superdex 75 10/300 GL column (GE Health Sciences), in which SCD1 with a detergent belt can be well separated from soluble cyt b_5_ and b_5_R. The FPLC buffer was prepared with deionized ultra-pure water, and O_2_ in buffer was removed by three cycles of purging with Argon gas before the addition of detergent. 1 mM EDTA was added and incubated with samples for 10 min before loading into FPLC. The peak fractions of SCD1 were concentrated to ∼50 μM. Protein concentrations were determined by a protein colorimetric assay (Bio-Rad). ∼200 μL of proteins samples together with flow-through buffers (during concentration) as blank controls were accurately weighed and sent for ICP-MS (Agilent 8800 Triple Quad ICP-MS) analyses at the Department of Earth & Atmospheric Sciences of University of Houston. All protein samples were digested in 2% HNO_3_. Calibrations with Fe, Zn, and Co standards were performed at the beginning and the end of each run. Metal contents of proteins were reported as the concentrations of metal in protein samples subtracted by those in the corresponding blank controls.

### UV-Vis spectroscopy and anaerobic titration

UV-Vis spectra were recorded using a Hewlett-Packard 8453 diode-array spectrophotometer (Palo Alto, CA). SCD1 solution in a tenometer was made anaerobic by 5 cycles of 30 s of vacuum followed by 4.5 min of saturating with argon. This anaerobic SCD1 solution was then titrated with anaerobic stock of dithionite in an air-tight syringe. The time courses of A_340_ in the reaction of H_2_O_2_ with SCD1 were recorded with an Applied Photophysics (Leatherhead, UK) model SX-18MV stopped-flow instrument. The observed rates, *k*_obs_, were obtained by fitting the time courses to biphasic exponential function. The fast spectral changes were monitored using the rapid-scan accessory with the stopped-flow machine and the optical species were resolved using the Pro-Kinetics program provided by Applied Photophysics.

### EPR spectroscopy

X-band EPR spectra were recorded with a Bruker EMX spectrometer (Billerica, MA) operating at 10 K. A Bruker dual mode resonator was used, and the parameters for the EPR measurements were: frequency, 9.58 GHz; microwave power, 4 mW; modulation frequency, 100 kHz; modulation amplitude, 10 G, and time constant, 0.33 s. The size of the EPR signals was based on peak-to-trough intensity. For power dependence study, the range of microwave power ranged from 200 mW to 0.1 mW, with a step of 3 dB. To prepare Fe(II)/Fe(II) SCD1 in EPR tube, resting SCD1 after EPR measurement was thawed and the atmosphere above the solution was flushed with N_2_ while small amount of dithionite was added to exhaust O_2_ in the solution. The capped EPR tube was then transferred into an anaerobic chamber, and 1 μM deazaflavin and 1 mM EDTA were added to the anaerobic SCD1 solution. The sample was then irradiated with white light for 15 min before being frozen for EPR measurement.

## Data availability

Data used for analyses are all included in the paper.

## Author contributions

J.S. conceived the project, conducted experiments, prepared figures, and wrote the paper. G.W. conceived the project, conducted experiments, and wrote the paper. B.S.P. helped the interpretation of EPR data and revised the paper. A.T. conceived and supervised the project and revised the paper. M.Z. conceived and supervised the project and wrote the paper.

## Competing interests

The authors declare no competing interests.

## Acknowledgements

This work was supported by grants from NIH (DK122784 to MZ and AT). We acknowledge V. Berka for his help with the dithionite titration, and G. Gerfen for insightful discussions on EPR data.

---

**Figure 1—figure supplement 1.**
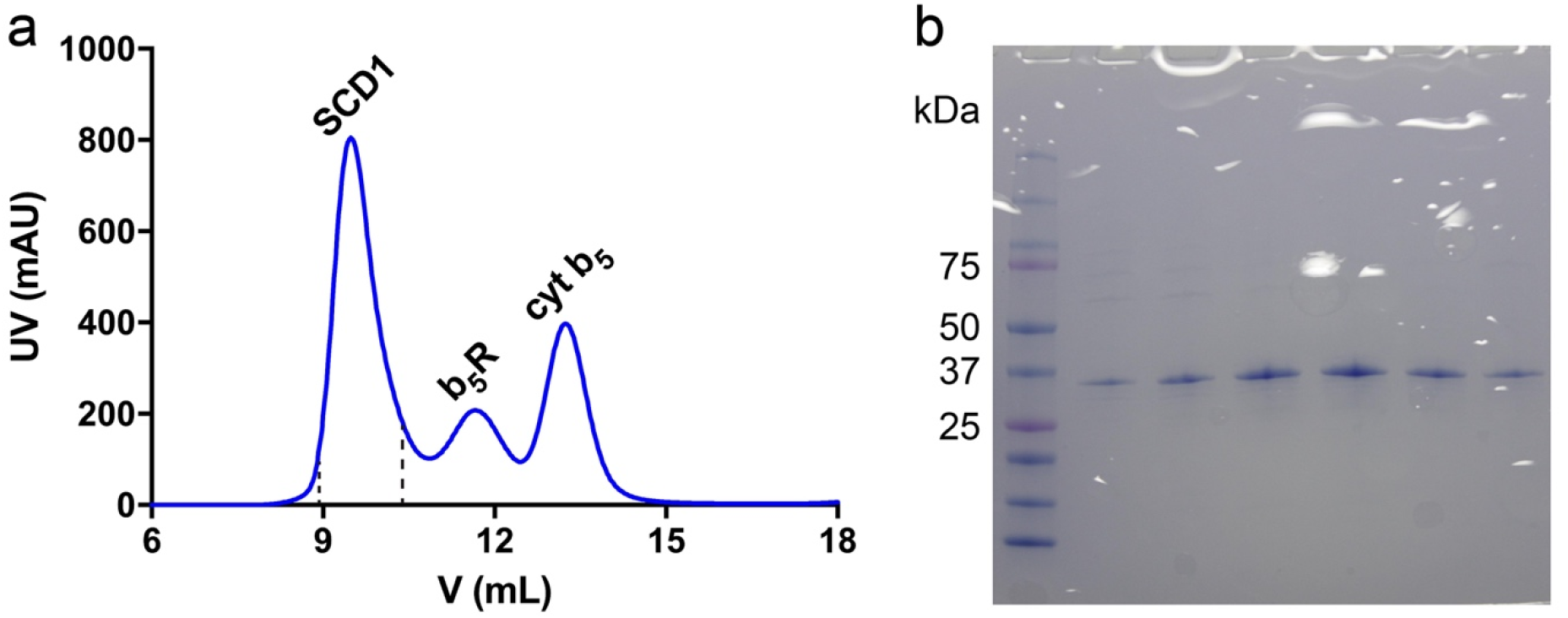
Purification of SCD1 after reaction. (**a**) SEC profile of reaction mixture containing SCD1, soluble b_5_R and cyt b_5_ in Superdex 75 column. Vertical black dashed lines indicate the region of elution collected for SCD1. (**b**) SDS-PAGE gel image of the collected fractions in (**a**).

**Figure 1—figure supplement 2.**
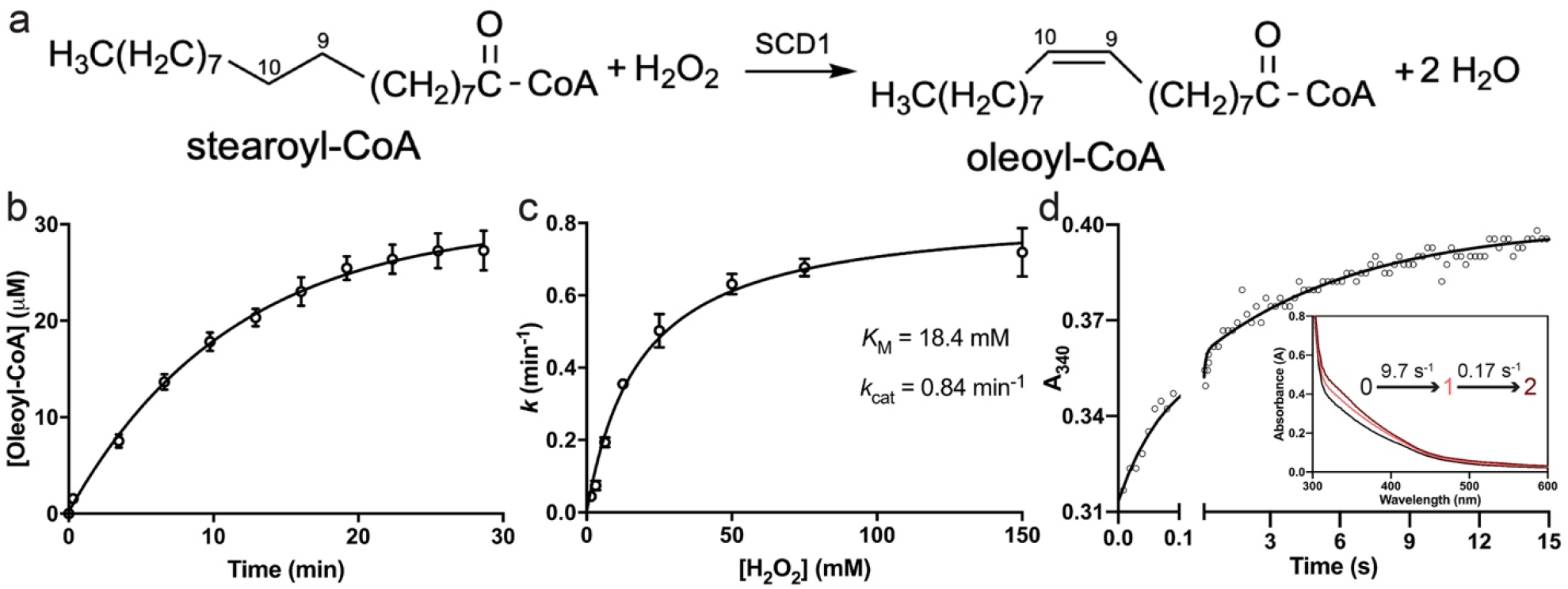
Reactivity of SCD1 in the peroxide-shunt pathway with H_2_O_2_. (**a**) Overall reaction scheme of the peroxide-shunt activity in SCD1. (**b**) Time course of oleoyl-CoA production with H_2_O_2_. (**c**) Michaelis-Menten kinetics of the peroxide-shunt pathway. (**d**) Pre-steady-state kinetics of oxidation of the diiron center in SCD1 by H_2_O_2_. The inset shows the spectra of three species (black, pink, and red) deconvoluted from kinetics analyses.

**Figure 2—figure supplement 1.**
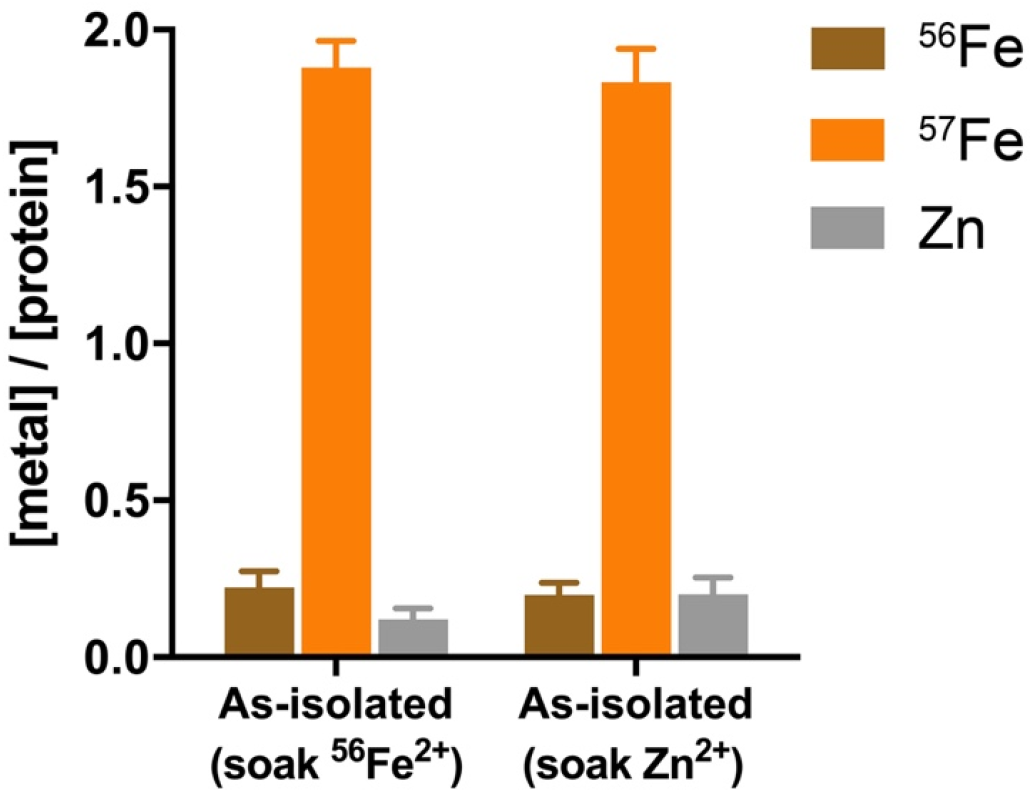
Metal content analysis of ^57^Fe-enriched SCD1 soaked with ^56^Fe^2+^ or Zn^2+^ without initiating reaction.

**Figure 3—figure supplement 1.**
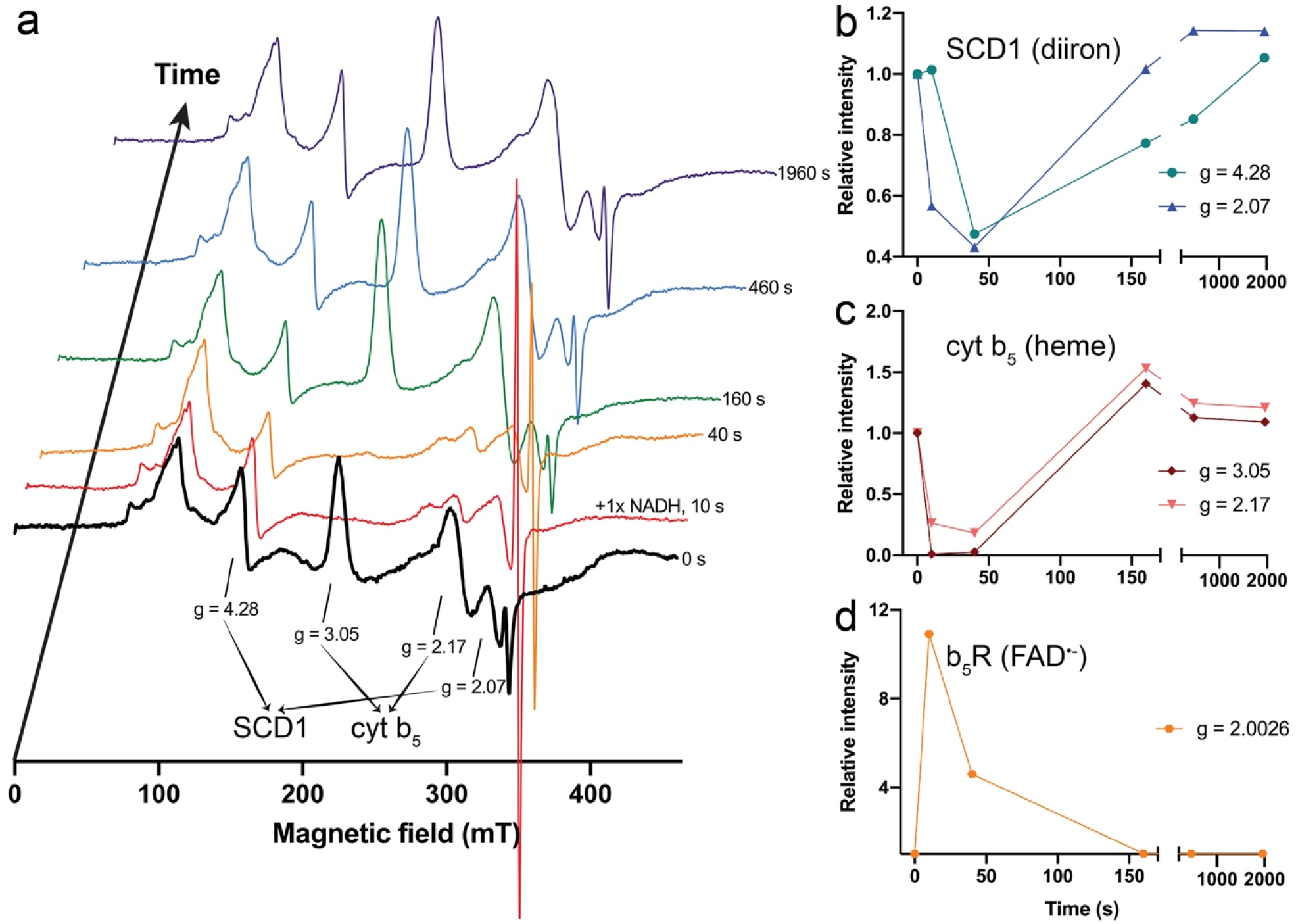
Redox cycle of SCD1-cyt b_5_-b_5_R complex followed by EPR spectroscopy. (**a**) EPR spectra of the ternary complex at 10 K. Characteristic *g*-values of resting state SCD1 and cyt b_5_ are labeled. Time stamps to the right of each spectrum represent the incubation time after the addition of one molar equivalent (1×) of NADH. The large sharp signal at *g* ∼2.0 popping up within 10 s is from the FAD^**-**^**·** in b_5_R. Time-dependent changes of relative intensities of the EPR species in: (**b**) SCD1 (*g* = 4.28 and 2.07); (**c**) cyt b_5_ (*g* = 3.05 and 2.17); and (**d**) FAD^**-**^**·** in b_5_R (*g* = 2.0026). The sharp signal at *g* ∼2.0 from SCD1 at resting state is not considered due to its significantly smaller size compared to the strong signal from FAD^**-**^**·**.

**Figure 4—figure supplement 1.**
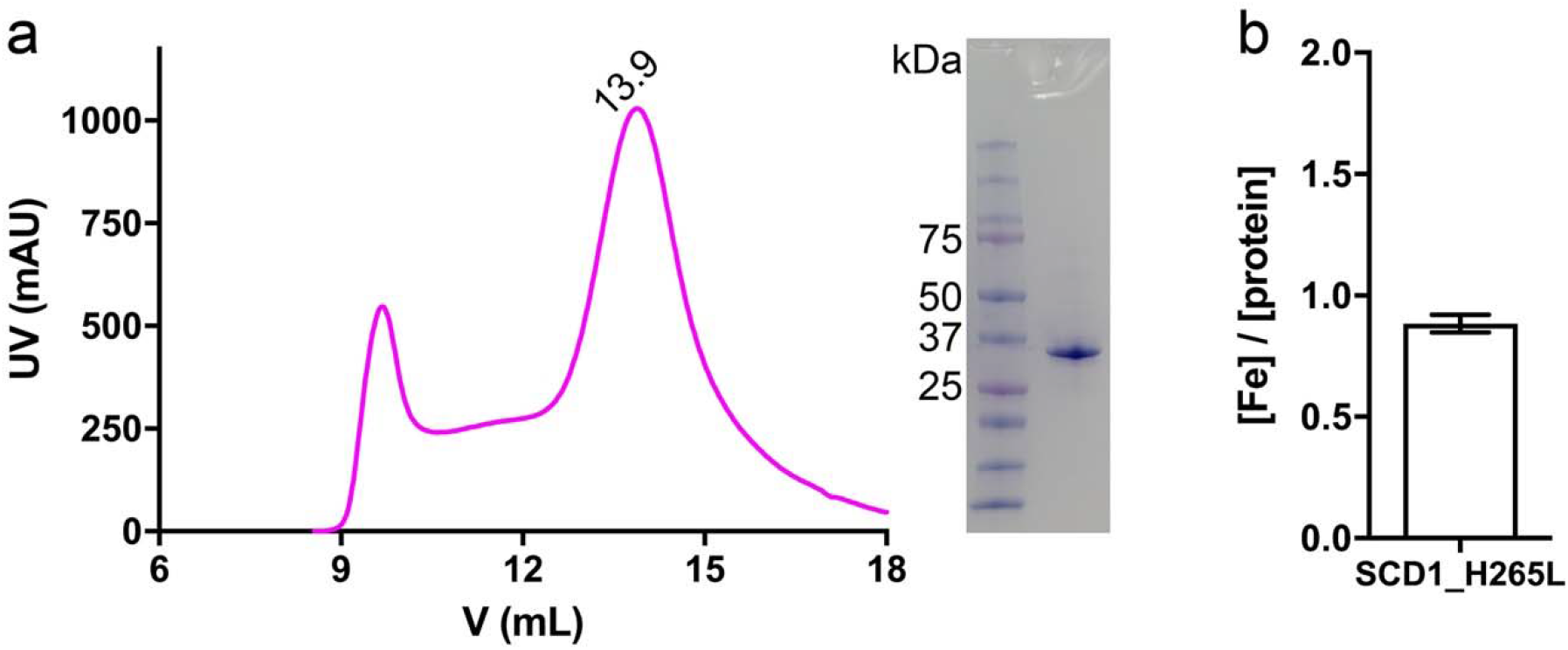
Purification of SCD1_H265L. (**a**) Left: SEC profile of SCD1_H265L in Superdex 200 column. Right: SDS-PAGE gel image of peak fractions. (**b**) Fe content of SCD1_H265L.

**Figure 4—figure supplement 2.**
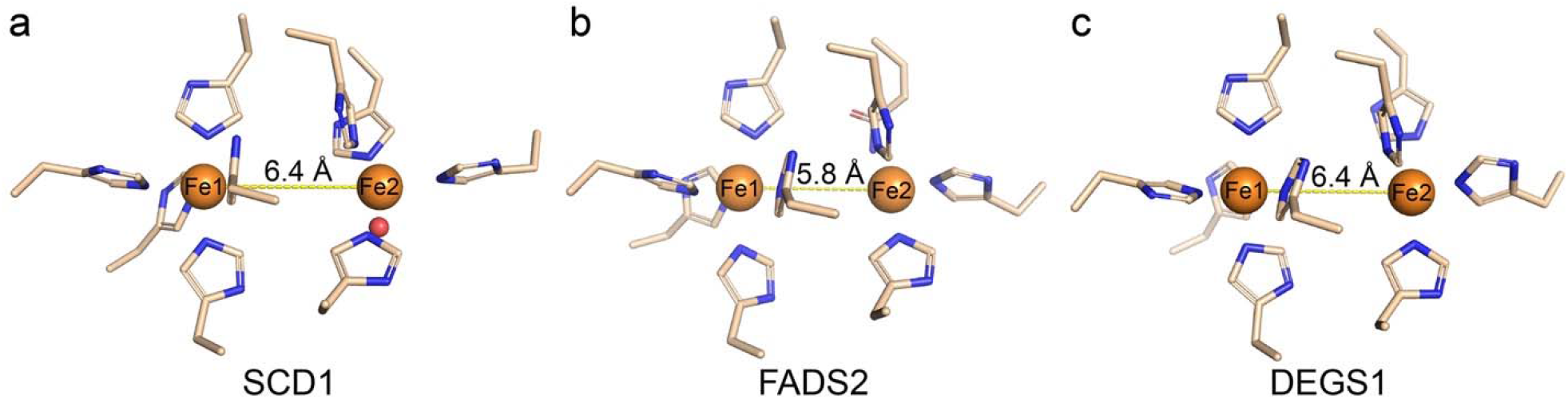
Comparison of diiron centers in some mammalian membrane-bound enzymes. (**a**) Crystal structure of the diiron center in mouse SCD1 (PDB ID: 6WF2). AlphaFold2-predicted models of active sites in: (**b**) human fatty acid desaturase 2 (FADS2, UniProt ID: <u>O95864</u>); (**c**) human sphingolipid Δ4-desaturase-1 (DES1, UniProt ID: <u>O15121</u>). Fe ions are placed at coordination distances to ligand residues in the predicted models.

## Notes

### Competing Interest Statement

The authors have declared no competing interest.

